# Selective and invariant features of neural response surfaces measured with principal curvature

**DOI:** 10.1101/2019.12.26.888933

**Authors:** James R. Golden, Kedarnath P. Vilankar, David J. Field

## Abstract

The responses of most visual cortical neurons are highly nonlinear functions of image stimuli. With the sparse coding network, a recurrent model of V1 computation, we apply techniques from differential geometry to these nonlinear responses and classify them as forms of selectivity or invariance. The selectivity and invariance of responses of individual neurons are quantified by measuring the principal curvatures of neural response surfaces in high-dimensional image space. An extended two-layer version of the network model that captures some properties of higher visual cortical areas is also characterized using this approach. We argue that this geometric view allows for the quantification of feature selectivity and invariance in network models in a way that provides insight into the computations necessary for object recognition.

## 1 Introduction

The linear systems approach to classifying and predicting the responses of visual cortical neurons yielded foundational insights into how the brain represents the visual world [Enroth-Cugell and Robson, 1966, Campbell and Robson, 1968, Blakemore and Campbell, 1969, De Valois et al., 1982]. The responses of V1 neurons have been characterized to a degree by responses to grating stimuli and the mapping of linear receptive fields. However, a family of nonlinear responses, including endstopping, position invariance, grating phase invariance and gain control [Albrecht et al., 2003], have also been observed, bringing into focus the limitations of the linear systems approach for V1 neurons. These nonlinear responses are quantified primarily as tuning curves for narrowly-parameterized stimuli; here, we use the tools of differential geometry to quantify simulated neural responses in a new way.

Endstopping and gain control can be broadly cast as selective nonlinearites, in contrast to the invariant or tolerant nonlinearites in responses to changing position or grating phase. Selective and tolerant nonlinearities play an important role in understanding the responses of individual neurons in V4 and IT [Rust and DiCarlo, 2012]. Experiments quantifying selectivity and invariance in high-order visual neurons Rust and DiCarlo [2012] use the responses to many different types of images, and similar approaches are used to characterize individual units in deep networks used for visual object identification [Goodfellow et al., 2009]. Selectivity is measured by the change in response to texture-scrambled images of objects, and invariance is measured by the responses to translated, scaled and rotated objects on the background. The untangling theory of object recognition [DiCarlo and Cox, 2007], in which the ventral stream untangles the the manifolds corresponding to views of different objects in pixel space, provides a mathematical formalism for interpreting selectivity and invariance. We provide evidence that selectivity and invariance correspond to negative and positive curvature, respectively, of the neuron’s response surface in image space [Berkes and Wiskott, 2006, Erhan et al., 2010, Tsai and Cox, 2015, Horwitz and Hass, 2012, Zhu and Rozell, 2013].

Zetzsche and Krieger [1999] and Olshausen and Field [2005] discussed the use of the geometry of a neuron’s response in high-dimensional image space as a method of characterization. We previously developed a quantitative method for measuring the non-linearity of the geometry of a neuron’s response in 2D subspaces [Golden et al., 2016]. Tsai and Cox [2015] developed a non-parametric method for finding selective and invariant responses of units in multilayer networks. Paiton et al. [2019] describe how the curved geometry of the sparse coding network can be used to resist adversarial attacks. Here, we present a simple but computationally intensive mathematical framework for making the notions of selectivity and invariance precise. We describe measurements of the local curvature of neural responses in image space for the sparse coding network [Olshausen and Field, 1996] and the two-layer variance components network [Karklin and Lewicki, 2003, 2005]. We measure curvature of iso-response surfaces in high dimensions using methods from differential geometry and carry out computations using automated differentiation (ADMAT 2.0 for Matlab, Cayuga Research). For a given neuron, the principal curvatures of the response manifold correspond to an eigenvector decomposition into a set of image features, where the eigenvalues indicate the magnitudes of curvatures, and the signs of the eigenvalues indicate selectivity (hyperbolic curvature) or invariance (spherical curvature).

Selective and tolerant/invariant responses have been defined by how much a neuron’s response decreases from a maximum with parametric changes to a stimulus. This definition has been necessarily limited in physiological experiments because of the limit on how many images can be presented (although see [Ponce et al., 2019]). For computational networks, it is possible to directly measure the geometry of the response surfaces with automatic differentiation, which allows for a precise quantification of the image features beyond the stimulus that maximally affects a neuron’s response. We believe that while this definition and computational method may be difficult to implement in physiology experiments, it will be useful in characterizing single unit and layer responses in deep networks.

## 2 Background

The sparse coding network, as a model that learns an efficient representation of image data and reproduces some of the nonlinear properties of V1 neurons, will be used to generate responses. The sparse coding network can be described by an energy equation, where an image *I* is represented by a set of neurons with basis functions in the dictionary Φ weighted by the vector *x*.

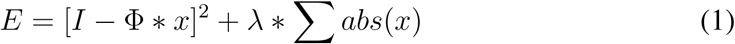

For a given Φ, x is found through gradient descent on this equation, subject to the squared error plus a cost function on the sparsity of the vector *x* (chosen in this instance to be the absolute value function), balanced by the coefficient *λ*. Φ is found over many iterations of alternating minimization of the energy equation to find *x* (the inner loop) and then to find Φ (the outer loop). In the following analysis of curvature, we use networks that have already been trained to have Φ that provides an accurate and sparse representation of natural image patches. We explore the curvature of the output responses of all the neurons in the network with respect to *x*, the image stimulus space.

The response manifold is the set of all possible responses for one neuron over image stimulus space. The principal curvatures of the neural response surface at a point in this space are a direct quantitative measurement of the high-dimensional features to which a neuron is selective and invariant. In Golden et al. [2016] and in Golden [2015], we interpreted illustrative examples of the sparse coding network in 2*D* that allow the responses to be visualized over the whole stimulus space and performed an analysis of the 2*D* subspaces of neural responses in *N* = 64 image space (Fig. 1). Here, we present measurements of the curvature of response surfaces in *N* = 64 image space.

**Figure 1:**
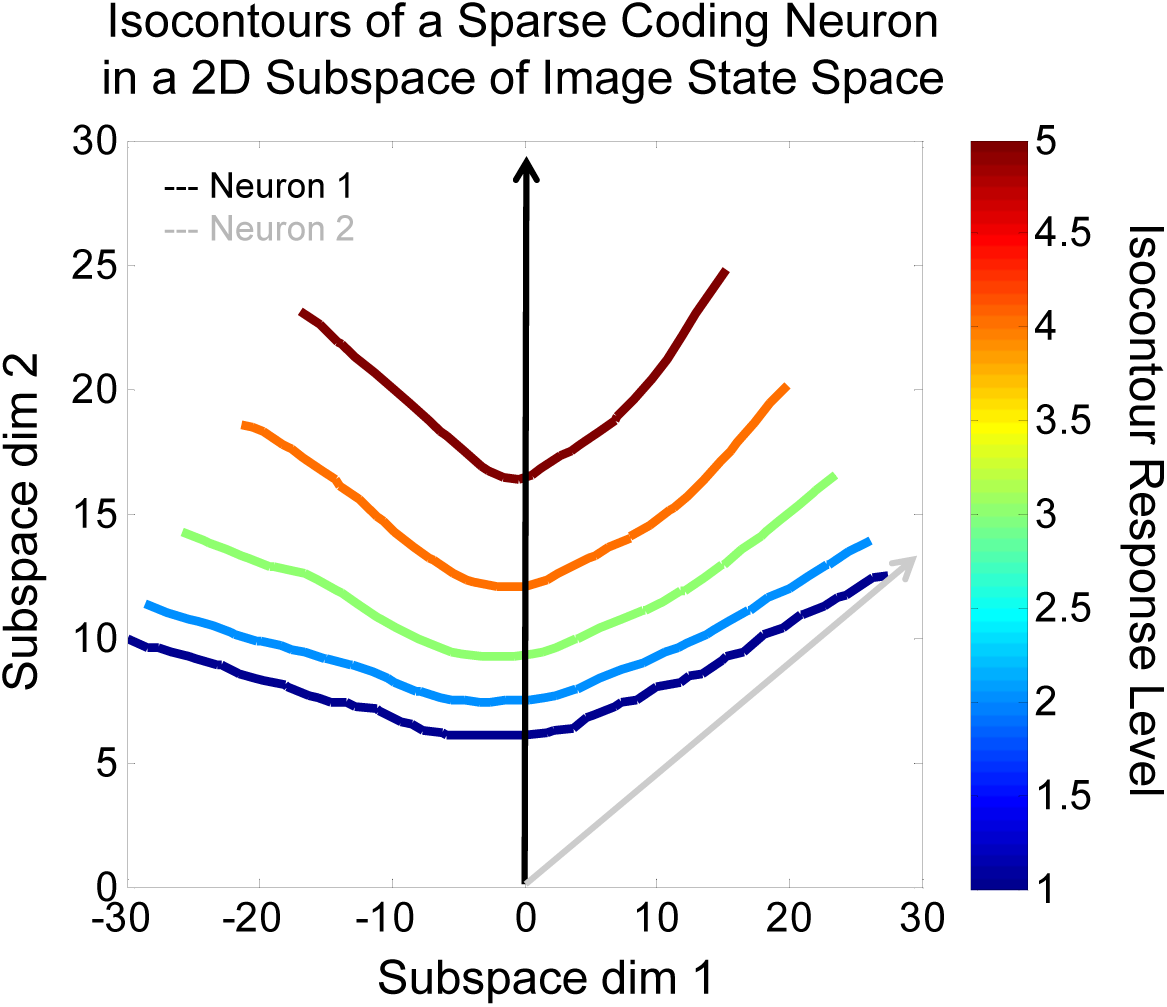
The iso-response contours for a neuron from a 6.4X overcomplete sparse coding network using a Gaussian cost function in a 2D subspace determined by this neuron’s basis vector (blue) and its closest neighbor (gray) at 60°. The basis vector points in the direction (0, 1), while the neighboring basis vector points at (0.9, 0.45). Note the curvature of the iso-response contours away from the neighboring basis vector.

The possibility that the sparse coding network could capture nonlinear effects like endstopping was mentioned in Olshausen and Field [2005] and thoroughly explored by Zhu and Rozell [2013], and the connection between endstopping and curved response manifolds was discussed by Field and Wu [2004]. We have provided evidence of the curvature of sparse coding response surfaces in low-dimensional subspaces, and we believe this allows for a new interpretation of how the sparse coding network finds a representation. The network provides an accepted method for placing encoding vectors to learn an efficient code that resemble V1 receptive fields. The recurrent inner loop step that finds an overcomplete sparse representation warps the encoding space according to the density of natural images in different regions of image space. A lattice representing image space becomes distorted and twisted when viewed in the representation space. The representation is made efficient by the placement of the encoding vectors that tessellate the space as well as how the space is warped by the encoding process. This distortion of the space is a way to view what the sparse coding network is doing from a more intuitive perspective, and it can be quantified with the curvature of the response surfaces.

The sparse coding network is a well-known model for network-level processing in V1, and serves as the basis for some of the deep learning networks that have become useful for object recognition [Le and Ng, 2013]. Measurements of the curvature of neural response surfaces from units in multi-layer networks will allow us to understand precisely the features to which units are selective and invariant.

We first discuss the output of applying the curvature measurement to the response surface of a neuron from the sparse coding network. We provide an interpretation according to feature selectivity and invariance. We discuss the important distinction between the curvature of the full response surface and an iso-response surface. Then we provide summary measurements of curvature over response surfaces and discuss comparisons between different measures and different networks.

## 3 Curvature in high dimensions

Curvature is a measure of how the surface normal changes locally in a quadratic fashion. For a surface in *N* -dimensional image space, the principal curvatures at a point are found by computing the inner product of elements of the Hessian (matrix of second derivatives) with the surface normal, followed by matrix division with the surface metric, and finally by the eigenvector decomposition of the result [Lee, 1997]. For a 2*D* surface in 3*D* space, the curvature equation is as follows:

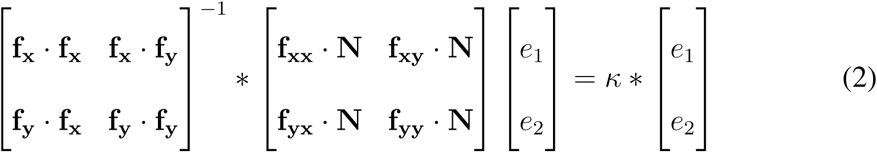

The principal curvatures consist of *N* eigenvectors and eigenvalues. For neural response surfaces, the principal curvatures are a quantitative measure of the features to which a neuron’s response is selective and invariant.

## 4 Curvature measurements in high dimensions

Consider a sphere in *N* = 64 dimensions with *R* = 4. Every normal section should show positive curvature equal to 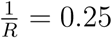. There are 64 principal curvature magnitudes and directions, with the magnitudes all equal to 1*/R* in the stem plot of 2 at left, and the directions in the 8×8 image patch plot at right. A principal direction in *N* = 64 space is a 64*D* vector, and can be shown as an 8×8-pixel image. When the curvature algorithm is applied to neural response surfaces, the images will allow some interpretation in the context of the basis functions of the network. The principal curvature computation results in the coordinate basis of the space as the set of principal directions. The eigenvectors of the Hessian can also be useful for interpreting the manifold curvature, but for a sphere these change at each point on the surface, whereas the curvature eigenvectors are invariant.

The measurement of curvature of the response surfaces of sparse coding neurons will quantify how nonlinear they are in high-dimensional image space. For the most part, predictions from the low-dimensional results of Golden et al. [2016] hold: curvature is a function of the sparsity, the angle between basis vectors and the overcompleteness of the network. The response surfaces in relevant subspaces are purely hyperbolic (all principal curvatures are negative), since the network is purely selective (although they are purely selective only at their maximum response value for an image of a particular contrast). We found that the response surfaces show the strong curvature in the directions of other neurons [Golden et al., 2016]. Specifically, the principal directions of the curvature of response surfaces ought to resemble the basis functions of other neurons in close angular proximity. The principal directions could have turned out to be any vectors in image space, but for high principal curvatures they are always similar to basis functions of other neurons in the network (it would be impossible for them to be identical, because the principal directions must be mutually orthogonal, and the basis functions are not in almost all cases). Since V2 neurons tend to be more nonlinear than V1 neurons, the response surfaces of neurons from the Karklin and Lewicki [2003, 2005] network show greater curvature than those of the sparse coding network, and that the maximum responses show positive curvature that is evidence of tolerance/invariance.

Since surface curvature is a local measurement, it will change as a function of position in image space. Due to the immense volume of *N* = 64 image space, we made measurements of the curvature at points of interest, and measured statistics of the principal curvatures over many such points. For example, if principal directions are the basis functions of nearby neurons, then perhaps the curvature magnitude is correlated with the angle between the surface’s basis function and other basis functions. As we have described before [Golden et al., 2016], we know the curvature of response surfaces is tied to the sparsity parameter *λ*, and therefore curvature will increase as the network is forced to be sparser. Along these same lines, curvature should also increase if there are simply more encoding vectors for the same space.

Another question is whether we will be able to observe global patterns in the curvature measurements (primarily in the principal directions). This seems somewhat unlikely, as the curvature should be more dependent on the point from which it is being measured. If a point on the neural response surface of neuron *A* is orthogonal to the basis function of neuron *B*, there should not be much curvature in the direction of neuron *B*; but if another point is only 20° away from neuron *B*, then there will likely be curvature at that point, in that direction. However, it may be possible to derive an approximate formula that predicts the curvature of a neuron at a particular point as a function of the basis set, which could allow for a feedforward approximation of the response of the sparse coding network. This approximation with quadratic basis functions would resemble the network that implements slow feature analysis [Wiskott and Sejnowski, 2002], although it would still be more complex as the curvature for neurons from the slow-feature analysis network is the same at every point.

Consider the output of the curvature measurement for a neuron in the sparse coding network shown in Fig. 3. In order to completely characterize a neuron with nonlinear behavior throughout the high-dimensional space, some form of analytic function or a lookup table would be necessary (although impossible, in the practical sense, in high dimensions). This curvature measurement of a response manifold yields only local information, but gives a measure of what the manifold is doing locally in the full high-dimensional space. The preceding claims about curvature of these manifolds can only be supported by numerically measuring curvature at many points on many response manifolds.

**Figure 2:**
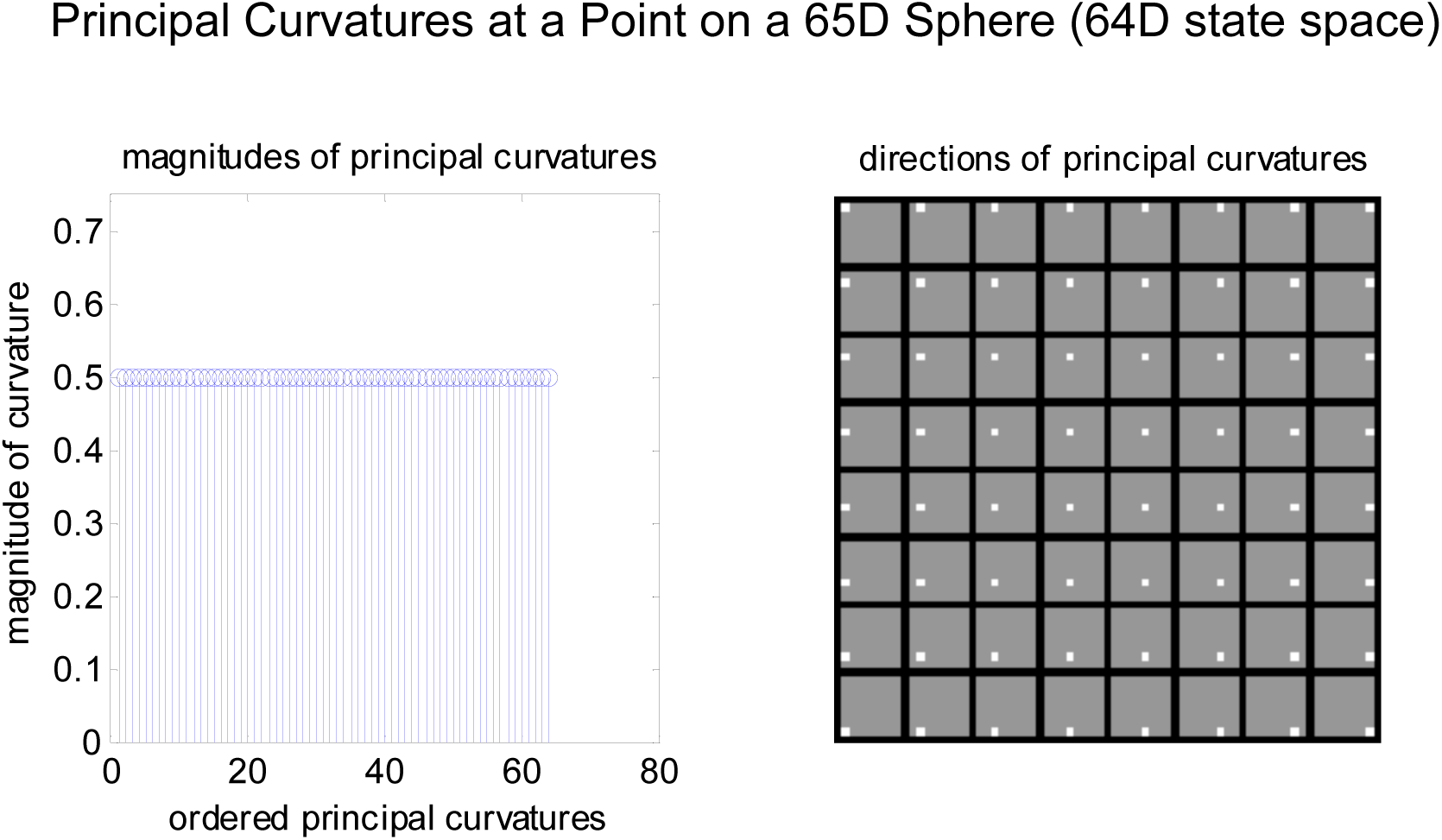
The principal curvatures and principal directions of the hypersphere with R = 2 in *N* = 64 image space. Note the curvature magnitudes are all equally 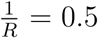, and the principal directions can be represented as 8×8-pixel images in the *N* = 64 space, here corresponding to the coordinate basis.

**Figure 3:**
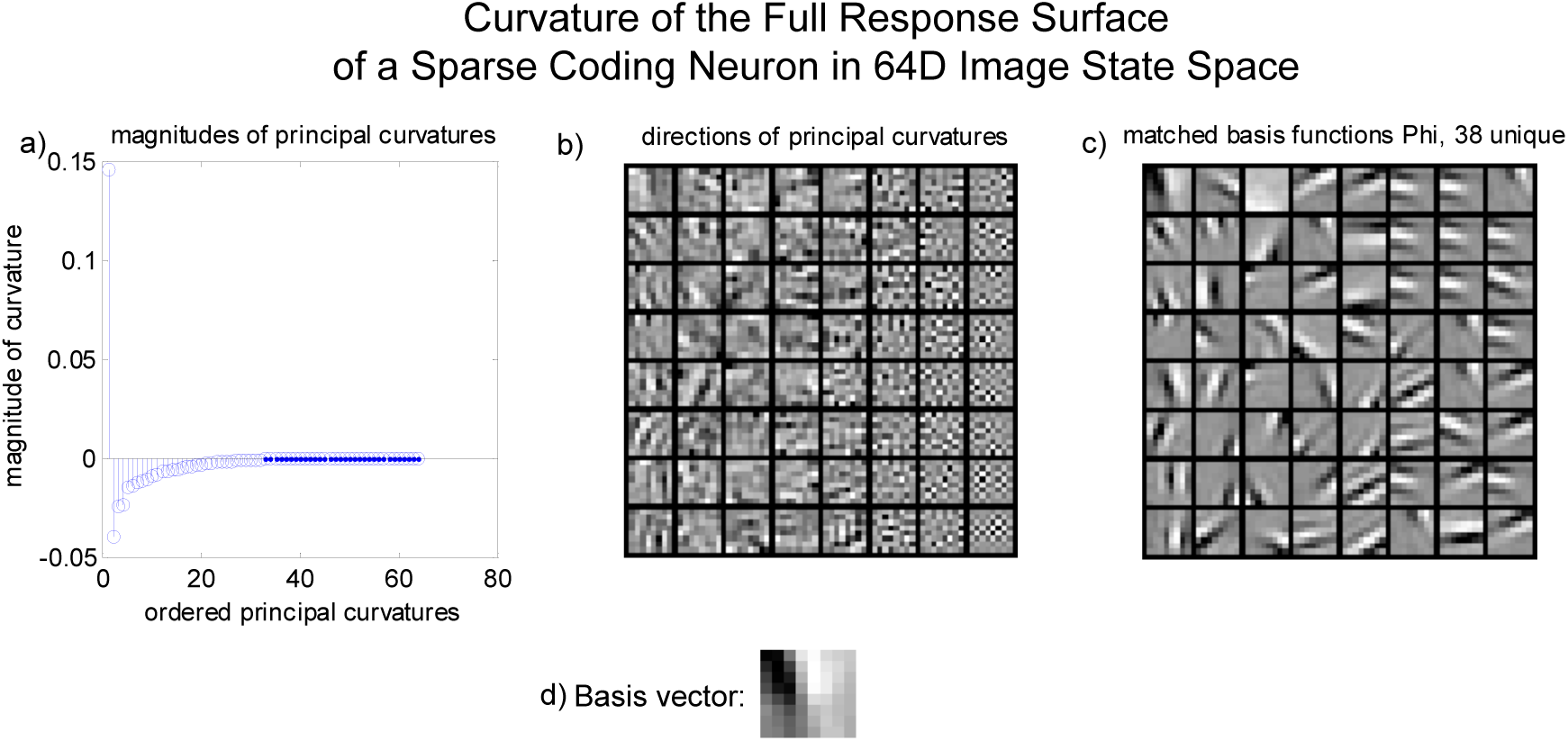
The curvature on the response manifold of one neuron at one point. a) The principal curvatures, which represent the magnitude of the curvature. b) The principal directions of the curvature. c) The closest basis function to each principal direction in terms of inner product. Note that the principal curvatures with the strongest magnitudes have directions that match reasonably well with basis functions, although they are not perfectly matched in terms of spatial frequency and phase.

Like the measurement for curvature at a point on a sphere in high dimensions above in Fig. 2, both the principal curvature magnitudes and directions are shown in Fig. 3. This measurement provides some support for the hypothesis that the directions of strongest curvature will be toward other basis vectors from the sparse coding network. As we have seen, the principal curvatures result from an eigenvector decomposition of product of elements of the Hessian with the surface normal followed by matrix division with the metric. The eigenvector decomposition is also used in principal components analysis (PCA). In PCA, the largest eigenvector is the direction of greatest variance in the data; the second-largest eigenvector is the direction of greatest variance in the data orthogonal to the first eigenvector, and so on. For the principal curvature eigenvectors, the largest is the direction of greatest curvature, the second largest is the direction of greatest curvature orthogonal to the first, and so on [Berkes and Wiskott, 2006]. The principal directions could have been any vector in *N* = 64 space for this particular response surface, but they are clearly related to the basis functions of other neurons in the network.

In other words, the directions of greatest curvature on the response surface are quite similar to the basis vectors of other neurons. The inhibition due to the nonlinear inner loop stage that finds the overcomplete sparse representation results in large degrees of curvature. To demonstrate this, Fig. 3 shows the basis functions that most closely correspond with each of the principal directions (in terms of largest inner product). There is a striking correspondence between the principal directions and the basis vectors of the network. The principal directions with the largest curvature correspond to basis functions that are less than 90° away from the basis vector representing the response surface. These relationships will be quantified in a statistical manner below. Interestingly, the principal directions are not the same as the basis functions. This may be because the principal directions are constrained to be orthogonal, and the basis functions for the overcomplete sparse networks are not orthogonal; however, this is also the case for the critically-sampled network that is not overcomplete. Alternatively, this is evidence that the principal directions yield additional insight into the highly curved manifold learned by each unit in image space.

One possible confounding problem with this analysis is that the largest curvature is in the direction of the neuron’s own basis vector. This is information about the contrast response of the neuron: when the magnitude of the stimulus is increased in the direction of the basis vector, the neuron’s response increases nonlinearly. As discussed earlier, the curvature of the iso-response surface of the neuron’s response manifold will not have this confounding problem, as the curvature will be measured with the basis vector as the surface normal. Below in Fig. 4 is the curvature analysis for an iso-response surface of the neuron [Bian et al., 2011, Chang et al., 2010]. This is quite similar to the curvature of the full surface, except there is no curvature in the direction of the encoding vector. The encoding vector is the normal vector to an isosurface, so the isosurface curvature measure serves as the more interesting characterization.

**Figure 4:**
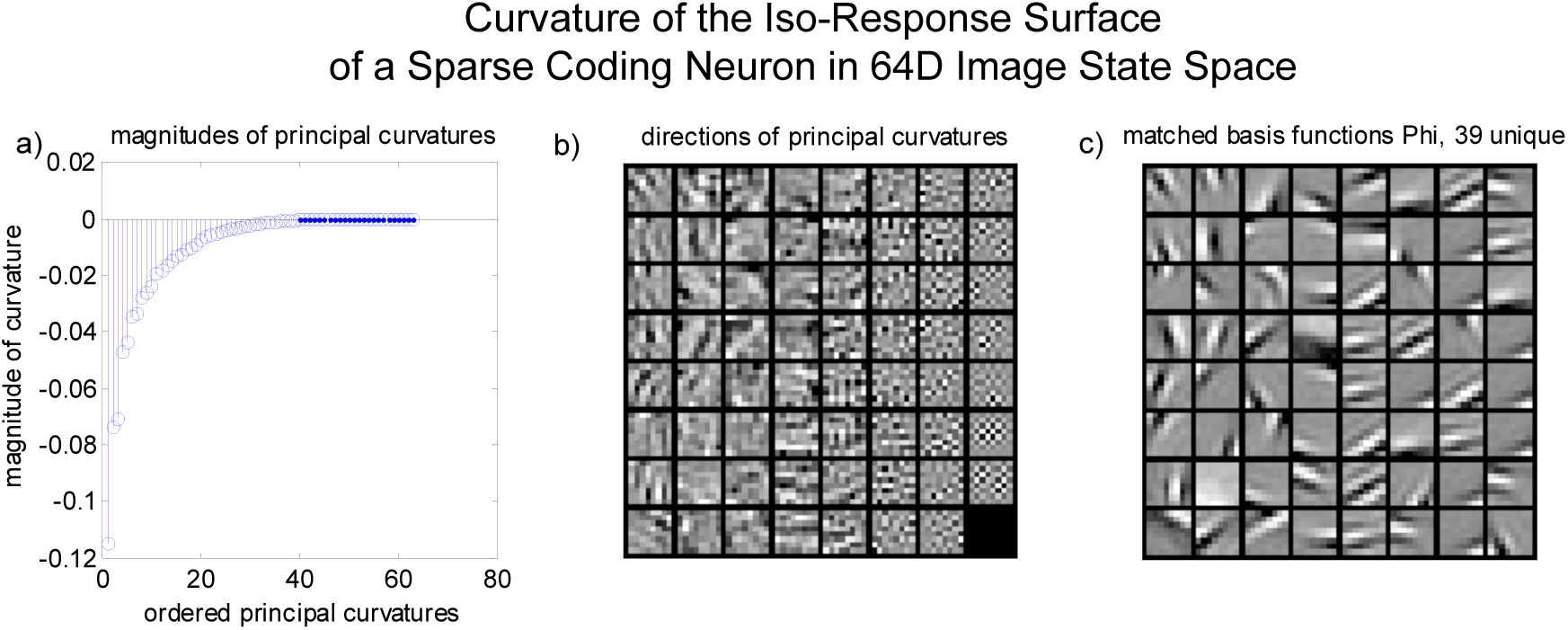
a) Curvature of the iso-response surface. b) The principal directions of the iso-response surface curvature do not include the neuron’s own basis vector (although they are (*N –* 1)-dimensional vectors). Therefore, the curvature of the iso-response surface is more informative about how the response manifold is warped by the presence of other neurons. For a neuron from the sparse coding network, at a point in image space identical with the neuron’s basis function, all of the principal curvatures of the isosurface are negative, indicating pure selectivity as well as the maximum value of the surface for a given contrast.

As for the magnitudes of the principal curvatures, the interpretation is not as obvious. First, note that the magnitudes of curvature are different for the iso-response surface and the full surface. This is in part an inherent difference in iso-surface curvature. Consider a sphere in *N* = 3 space, *x*^2^ + *y*^2^ + *z*^2^ = *R*^2^, with iso-response surfaces that are circles. For a constant *z* value, the radius of the iso-response circle is defined by a choice of the point on the sphere. When *x*^2^ + *y*^2^ is small, *z* is large, and the iso-response circle has a small radius and therefore a large curvature. When *x*^2^ + *y*^2^ is large, the radius of the iso-response circle approaches the radius of the sphere. The curvature magnitude of any iso-response surface therefore has a lower bound defined by the radius of the full surface. Iso-response curvature magnitudes should not completely match curvature magnitudes of the full surface. As long as isosurface curvatures are only compared to other isosurface curvatures, the comparisons will be meaningful.

Here we will also consider the response surface curvature of neurons from the second layer of the Karklin & Lewicki network. Based on Rust and DiCarlo [2012], neurons upstream in the ventral pathway will likely show both increased selectivity and increased invariance. The second-layer neurons of the Karklin & Lewicki network have been shown to capture aspects of invariance that are not seen in sparse coding neurons. The curvature of their response manifolds should therefore be greater than the curvature of selectivity in V1-like sparse coding neurons, and this is the case as shown below in Fig. 5. There are clearly strong positive and negative curvatures, and the absolute magnitudes are much greater than what is seen in sparse coding responses. Finally, note the invariant responses to vertically-oriented features, while selective responses are due to horizontally-oriented features.

**Figure 5:**
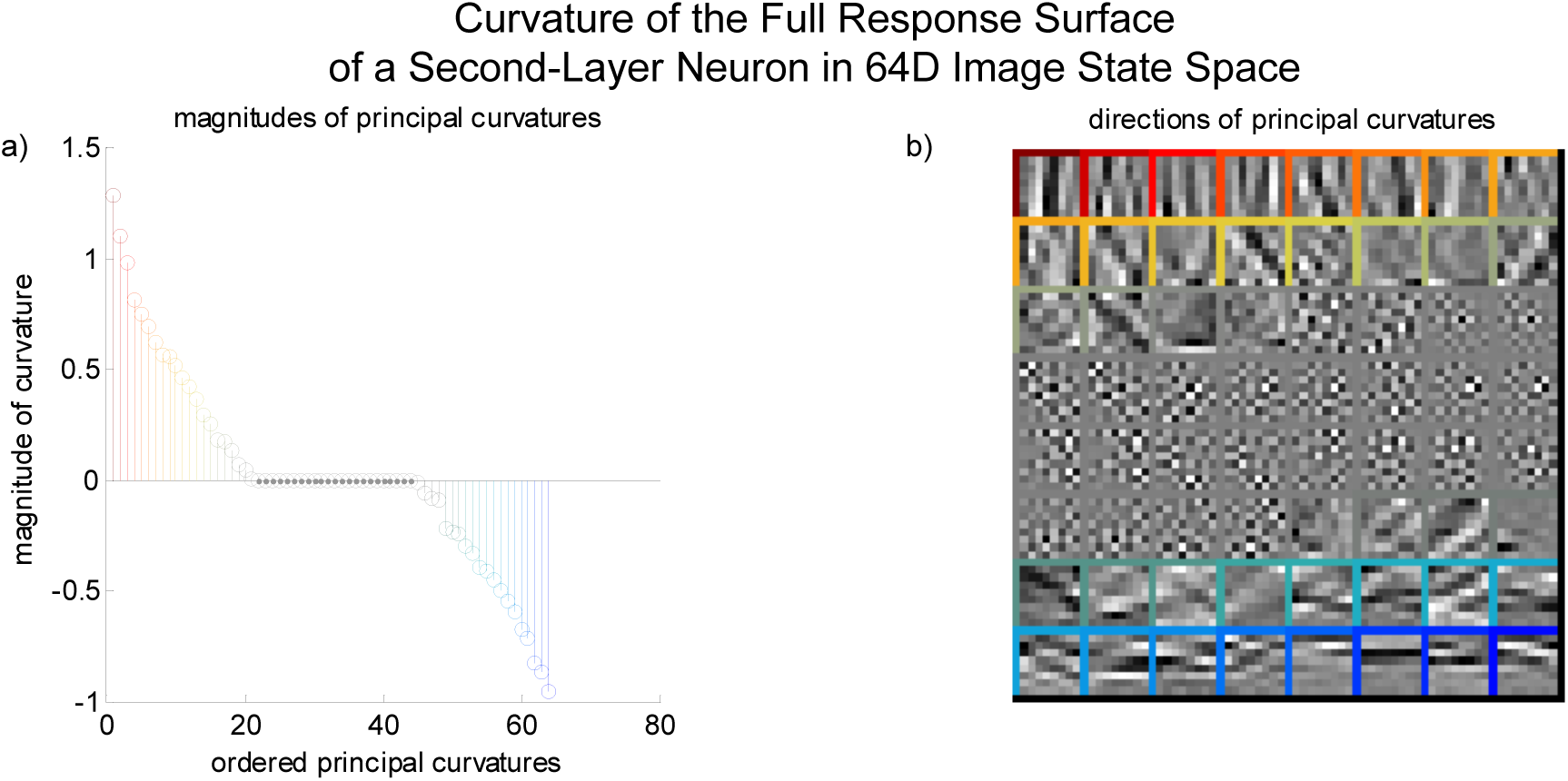
The curvature of a second-layer neuron from the Karklin & Lewicki network. Note the curvature is several orders of magnitude larger than that of the sparse coding neuron above in Fig. 3 and that there are strong positive and negative components. The neuron is invariant toward vertically-oriented features and selective for horizontally-oriented features.

The measurement of the curvature of a neuron’s response in this full high-dimensional space provides support for the idea that selectivity is identical to negative curvature of the response manifold. By making numerical measurements of the magnitude of the selectivity and the directions in which the neuron is selective, we have a new way of conceptualizing a neuron’s selectivity. Along with that comes an interpretation of the curvature, such that we can use the method to describe exactly how selective a neuron is to a set of specific image features. These measurements may not be feasible for physiological experiments because they are reliant on equations describing a response, but it may be possible to use them to make predictions from a model that captures aspects of the responses of cortical neurons. For example, if we know a neuron’s receptive field, and the receptive fields of others nearby in image space [Tsai and Cox, 2015], then we ought to be able to simulate responses using the sparse coding network to make predictions about where the isocontours will fall in the image space. Further, it could be useful in interpreting deep networks, where it can be difficult to understand the response properties of single units. Simple ways of testing feature selectivity and invariance based on a network finding the representations for a large batch of images [Goodfellow et al., 2009] may be compared with this theoretical measurement of selectivity and invariance.

## 5 Summary Measures of Neural Response Curvature

We collected summary statistics on the curvature of neural response surfaces from the sparse coding network at points in image space. The response surfaces could be approximated as functions of some high-order polynomial, so the curvature, which is a second-order measure, changes at every point. Since the surfaces are high-dimensional objects, it is not practical to measure or parameterize the local curvature over the whole image space. We chose to measure the curvature at a subset of the images of a constant contrast (equivalently, the surface of a sphere in image space). For one category, we chose sample points within 90° of each basis vector, because we have found that there is strong curvature at those points of response surfaces. In addition to the points in image space near the basis vectors, we have also measured curvature at natural image points from the training set. This allows us to measure the response surface curvature for points of image space that the network actually visits when it is encoding real natural image data. For comparison, we have also examined the curvature at random points in image space. A further complexity is that we are interested in the curvature of the iso-response surfaces. Measuring the curvature at a large number of sample points throughout image space allows us to look at summary statistics, like how many eigenvectors show non-zero curvature and how much of the curvature is concentrated in some number of the eigenvectors. We can compare these summary measures to each other for curvature points near the basis vectors, curvature at natural image points and curvature of the iso-response surfaces at both of these sets of points. These measurements were also carried out for the second-layer neurons from the Karklin & Lewicki network.

A simple summary of the curvature measure over many points is the distribution of the principal curvature values. Since the *N* = 64 space has been reduced to *N* = 40 by whitening (to speed up the learning of basis functions), 24 principal curvatures will be zero. For each point at which the curvature is measured, there will be 40 values contributing to the distribution. Fig 6 shows that the curvature of the iso-response surfaces measured at the basis vector image were nearly all negative, indicative of pure selectivity. The magnitude of every positive value was less than 0.001 at a radius of 1, so the positive curvature was extremely small. The sparse coding network therefore produces neurons with almost pure selectivity.

**Figure 6:**
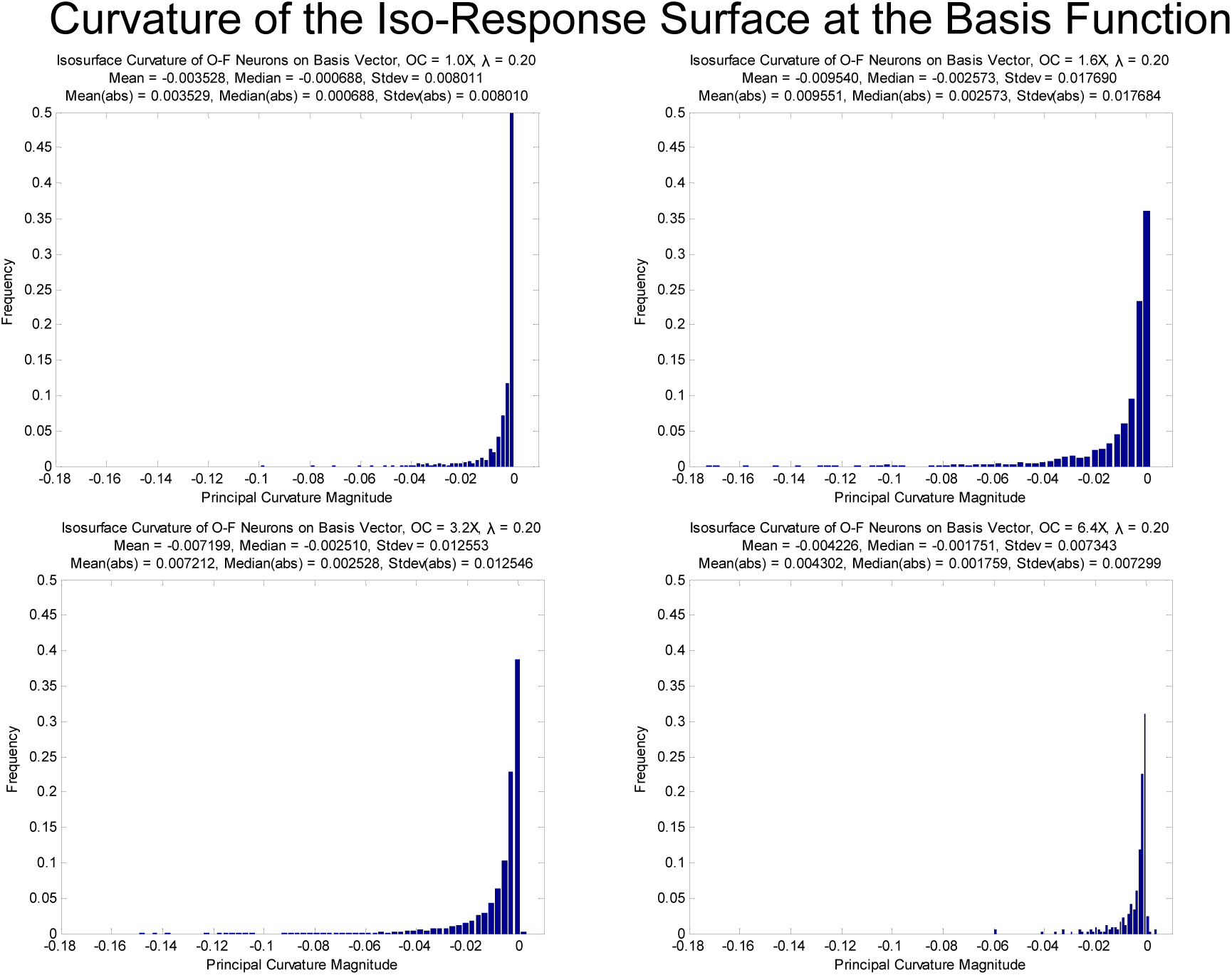
The distribution of principal curvature magnitudes for points on the basis vectors on the iso-response surfaces of neurons from the Olshausen & Field network at four degrees of overcompleteness.

A simple comparison to make is between the curvatures of response surfaces for the Olshausen & Field network in Fig. 7a with the second-layer neurons from the Karklin & Lewicki network in Fig. 7b. The immediate difference is simply the magnitude of the distributions (note the different scales on both axes). The principal curvatures for the second-layer neurons are two orders of magnitude greater than those that are found in the single-layer network.

**Figure 7:**
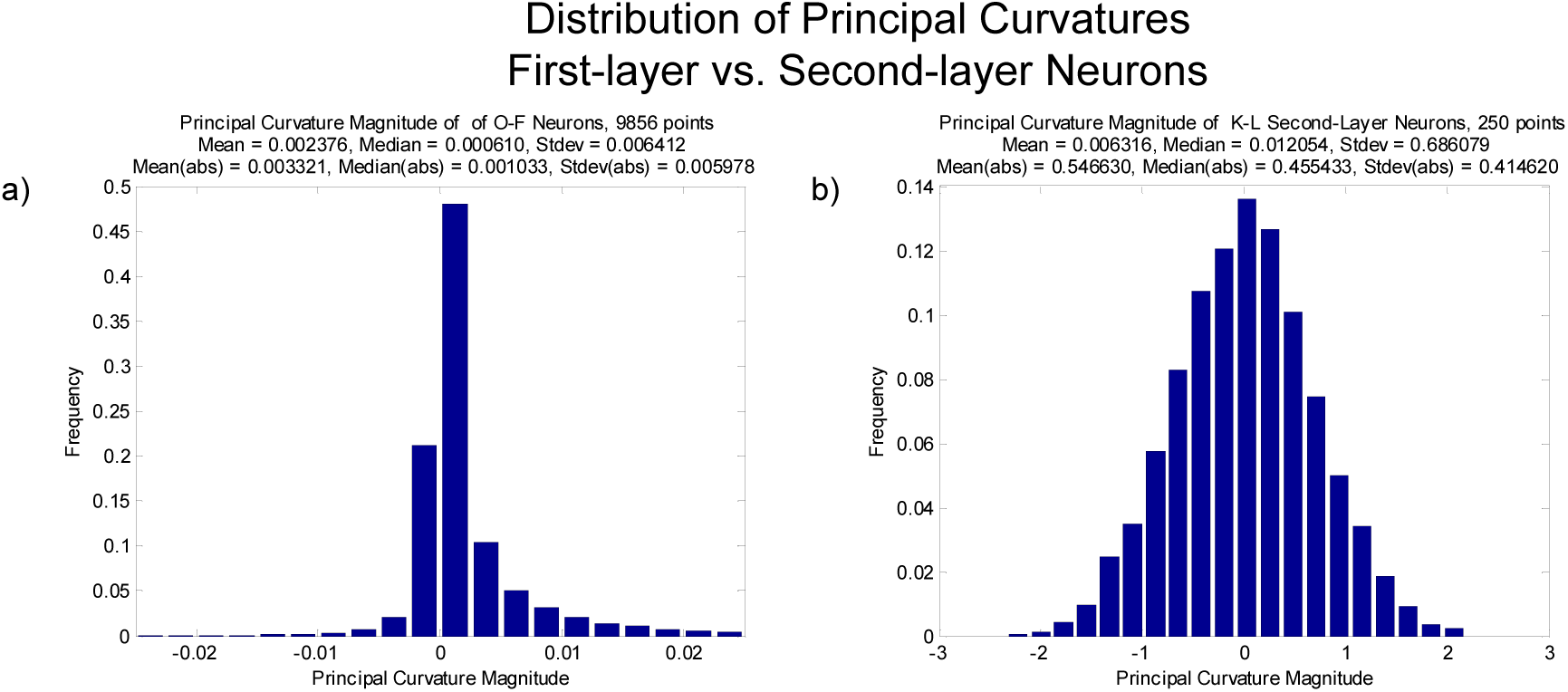
a) The distribution of principal curvature magnitudes for points near the basis vectors on the full response surfaces of neurons from the Olshausen & Field network. b) The distribution for second-layer neurons from the Karklin & Lewicki network. The median value is two orders of magnitude larger than the median value in a).

From an example of the curvature at a single point on a neural response surface, it is clear that a large percentage of the total mean curvature is concentrated in a small number of principal curvatures. In order to quantify this, we calculated the number of principal curvatures/eigenvalues which account for 95% of the total curvature as a measure of the degree of nonlinearity of the response surfaces. For 8×8-pixel natural image patches, the whitened space is 40*D*, and the mean and median number of principal curvatures that account for 95% of the total curvature at 10,000 image points are both about 18 dimensions, shown in Fig. 8. This means that 95% of the principal curvature is concentrated in less than half of the dimensionality of the response surface. For second-layer Karklin & Lewicki neurons, the median value is 31 dimensions, so the curvature of these neurons is spread out over more of image space than the single-layer neurons.

**Figure 8:**
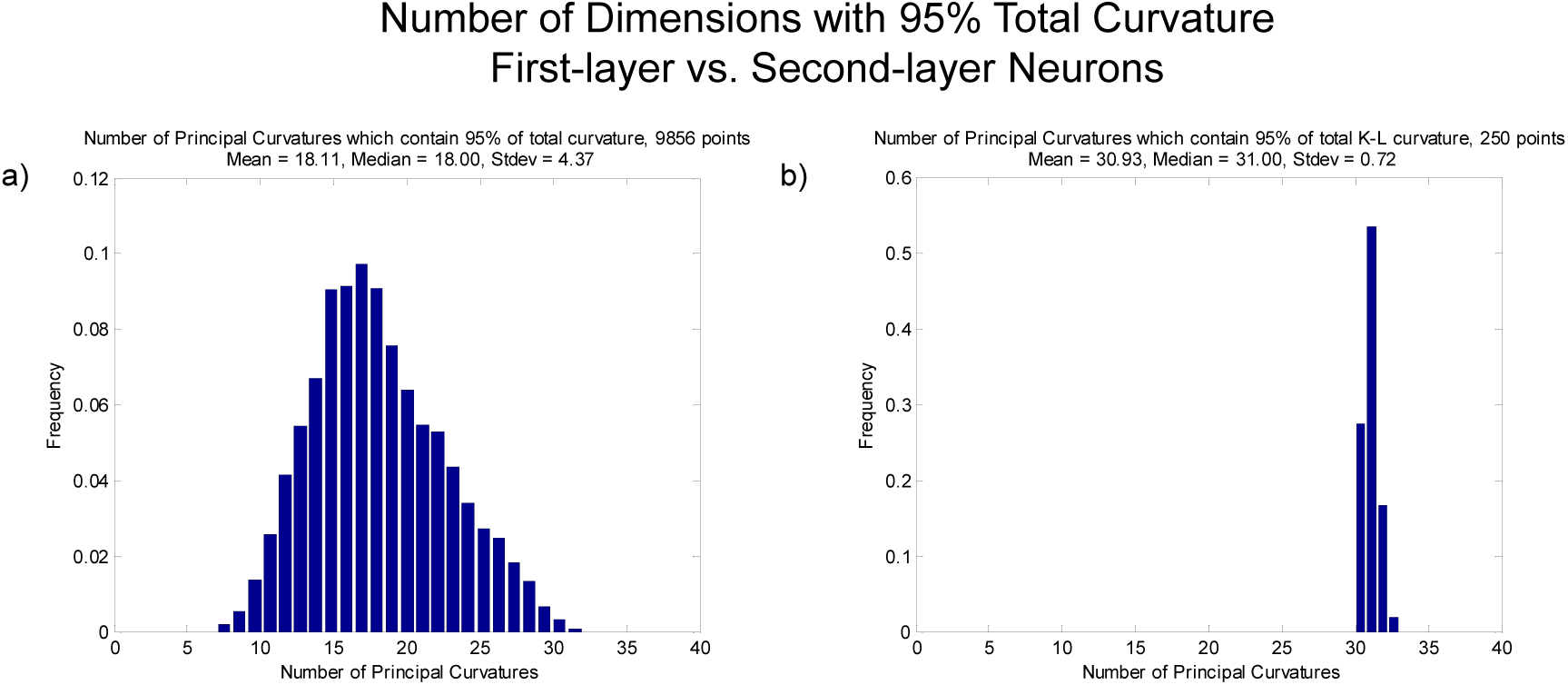
A histogram of the number of principal curvatures/eigenvalues for isosurfaces which account for 95% of the total absolute curvature. a) Olshausen & Field neurons, with a median of 18 dimensions. b) Karklin & Lewicki second-layer neurons, with a median value of 31 dimensions.

In Fig. 9, we present a quantification of the effect of both overcompleteness and the value of *λ*. We measured the curvature of the isoresponse surfaces of neurons from the Olshausen & Field network at 1000 natural image points for different values of overcompleteness and *λ*. Fig. 9 shows plots of the mean curvature (the average of the principle curvature magnitudes) as a function of the angle between the basis vector and the natural image point at which curvature was measured. Generally, the closer the natural image is to the basis vector, the higher the curvature of the isoresponse surface of that basis function. When the image point is 90° from the basis vector, the curvature is usually zero. Finally, the curvature generally increases with overcompleteness and with the value of *λ*.

**Figure 9:**
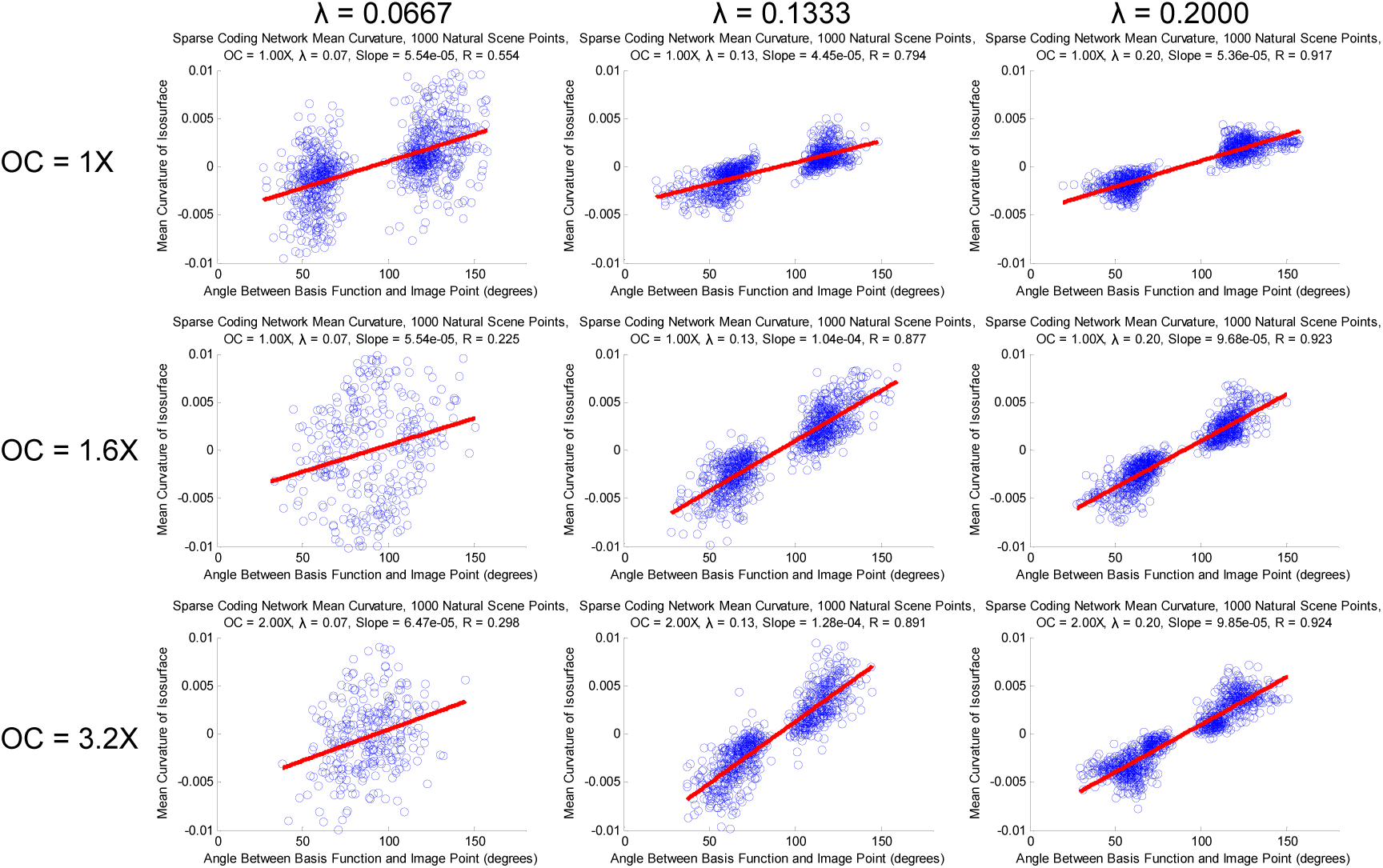
Plots of the mean curvature for isosreponse surfaces of Olshausen & Field neurons at natural scene points. Each plot is for response surfaces from a network trained at a certain degree of overcompleteness and a particular lambda value. The mean curvature is the average of the principal curvatures, and it is plotted as a function of the angle between the basis vector and the natural image point at which the curvature of the response surface was measured. Note that the mean curvature is about zero at 90°, and increases most dramatically for high overcompleteness and high *λ*. These plots demonstrate that mean curvature is a function of angle, sparsity and overcompleteness.

The plots of Fig. 9 can be summarized by the slope of the fit from each. The slope relates the angle between neurons to the curvature of the response surfaces, and the slope is clearly a function of both the sparsity (*λ*) and the overcompleteness of the network.

## 6 Conclusion

In expanding the measurements of Golden et al. [2016] into the full high-dimensional image space, we have considered how the curvature in the image space provides a direct quantification of the selectivity and invariance of a neuron’s response, and observed these measurements for a number of neurons. The increased increased curvature and feature selectivity of the response surface of neurons from the second layer of the Karklin & Lewicki network was demonstrated as compared to neurons from the sparse coding network. We measured summary statistics and showed that most of the curvature of a sparse coding neuron’s response is concentrated in 18 of 40 possible dimensions on average, while it is concentrated in 31 dimensions for second layer neurons. We quantified the effects of sparsity and overcompleteness on the curvature of response surfaces, and showed that both generally increase curvature as a function of angle. This new measure of selectivity and invariance provides a window into the features that drive a neuron’s response, both in the sparse coding network and a multi-layer network with more complexity.

## 7 Supplementary Material

**Figure 10:**
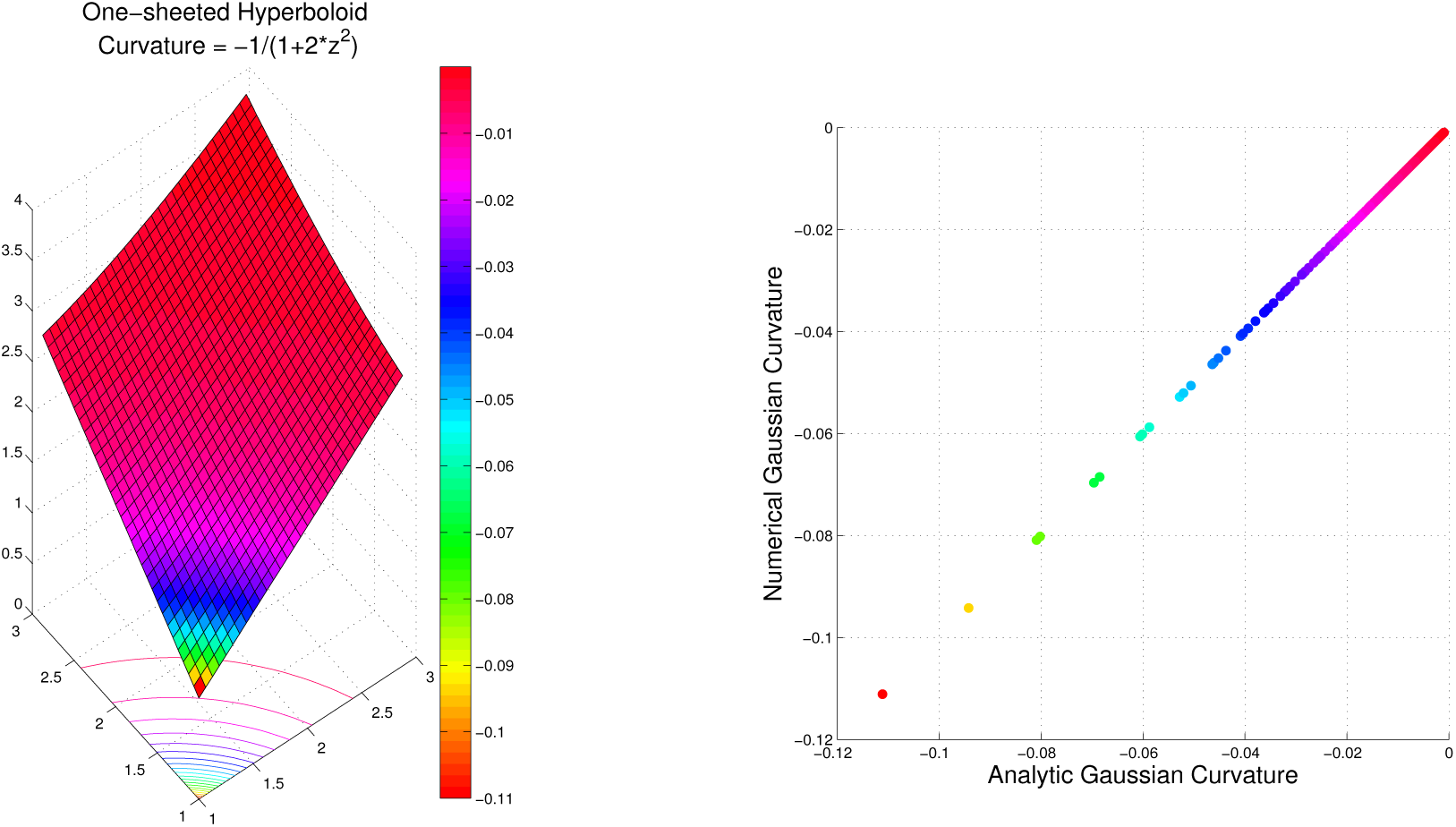
For *x*^2^ + *y*^2^ – *z*^2^ = 1, we show the surface with the numerical Gaussian curvature (the product of both principal curvature magnitudes) on the left, and a plot comparing the numerical curvature and the analytically determined Gaussian curvature. Note the perfect correspondence, supporting the accuracy of the numerical measurement process [Weisstein, 2001b,a]

